# Structure of the hexameric fungal plasma membrane proton pump in its auto-inhibited state

**DOI:** 10.1101/2021.04.30.442159

**Authors:** Sabine Heit, Maxwell M.G. Geurts, Bonnie J. Murphy, Robin A. Corey, Deryck J. Mills, Werner Kühlbrandt, Maike Bublitz

**Affiliations:** Department of Biochemistry, University of Oxford, South Parks Road, Oxford OX1 3QU, United Kingdom; Max Planck Institute of Biophysics, Max-von-Laue-Str. 3, 60438 Frankfurt am Main, Germany

## Abstract

The fungal plasma membrane H^+^-ATPase Pma1 is a vital enzyme, generating a proton-motive force that drives the import of essential nutrients. Auto-inhibited Pma1 hexamers in starving fungi are activated by glucose signalling resulting in phosphorylation of the auto-inhibitory domain. As related P-type ATPases are not known to oligomerise, the physiological relevance of Pma1 hexamers remains unknown. We have determined the structure of hexameric Pma1 from *Neurospora crassa* by cryo-EM at 3.3 Å resolution, elucidating the molecular basis for hexamer formation and auto-inhibition, and providing a basis for structure-based drug development. Coarse-grained molecular dynamics simulations in a lipid bilayer suggest lipid-mediated contacts between monomers and a substantial protein-induced membrane deformation that could act as a proton-attracting funnel.

## Main Text

The fungal plasma membrane H^+^-ATPase Pma1 regulates intracellular pH and generates a proton gradient that drives the import of nutrients^1^. The membrane potential generated by Pma1 can reach hundreds of millivolts^2^, requiring tight coupling of ATP hydrolysis to unidirectional H^+^-transport. Considerable effort has gone into characterizing Pma1^1^, not least because the enzyme represents a valuable drug target against severe invasive mycoses, particularly in immuno-compromised patients^3–4^. However, an 8 Å map from 2D electron crystallography^5^ is currently the highest-resolution structural information available for Pma1, hampering structure-based drug development.

Unlike any other known P-type ATPase, Pma1 forms hexamers that localise to specific ordered microdomains in the plasma membrane^6^. It is highly abundant and can form dense, paracrystalline arrays in starving or stressed cells^7,8^. In addition to the typical P-type ATPase core architecture, Pma1 carries auto-regulatory sequence extensions at both its termini ^9,10^ (Fig. S1a). The N-terminal extension is poorly characterised, but the C-terminal R domain has been postulated to auto-inhibit Pma1 in starving cells by immobilising the cytosolic domains^7^. Upon glucose sensing, Pma1 is activated by phosphorylation of the R domain.

Yeast studies have demonstrated that hexamer formation depends on association with lipids in the endoplasmic reticulum, which mediates surface transport via the secretory pathway^13–15^. Very-long-chain sphingolipids play a crucial role in Pma1 biosynthesis and targeting^13^, and Pma1 activity *in vitro* is stimulated by anionic lipids^14–15^. However, structural evidence to understand these findings has been lacking.

We collected single-particle cryo-EM data of hexameric Pma1 from *Neurospora crassa* in its auto-inhibited state and obtained a hexamer map at 3.3 Å resolution (Fig. 1a,b). An improved map for the monomer at 3.1 Å resolution was derived from the same data by symmetry expansion of the particles followed by focused 3D classification and refinement using a monomer mask (Fig. 1c, Fig. S2, Table S1). A monomer model was built and then extended into a hexamer by symmetry-assisted placement of five additional copies into the hexamer map.

**Figure 1.**
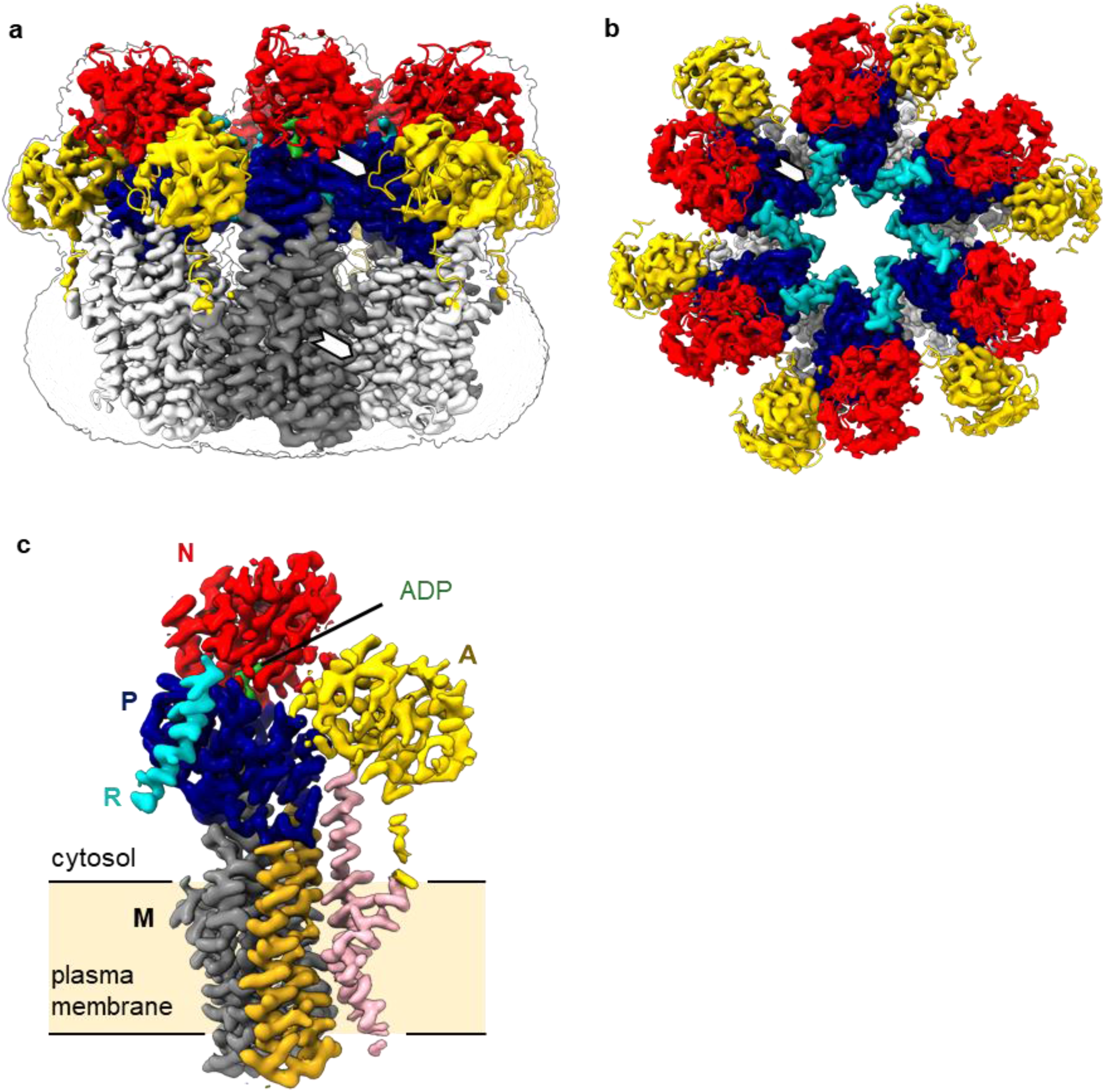
The cryo-EM structure of monomeric and hexameric Pma1. **a,** Overlay of cryo-EM map and structural model of the Pma1 hexamer in side view (with map outline delineating the detergent micelle). Nucleotide-binding (N) domain in red, actuator (A) domain in yellow, phosphorylation (P) domain in blue, regulatory (R) domain in cyan and ADP in green. The M domains of individual monomers are shown in alternating shades of grey. White arrows indicate contact sites between monomers **b,** Top view of the hexamer. **c,** Cryo-EM map of the Pma1 monomer subunit. Colouring as in a, except the ten helices of the membrane domain (M) are colored in pink (M1-2), gold (M3-4) and grey (M5-10).

Pma1 has a typical P-type ATPase fold: the M domain comprising the membrane-spanning α-helices M1 to M10, and the three cytosolic domains A, N, and P (Fig. 1c, Fig. S1a). The C-terminal auto-inhibitory R domain forms an α-helix that folds against the P domain. The model comprises >90% of all residues, lacking only the N-terminal extension, and two loops (A-M3 and M10-R), for which continuous density was lacking, probably due to flexibility.

The Pma1 hexamer forms a symmetric ring with a maximum outer diameter of 167 Å, and an inner cavity measuring 24 Å and 55 Å in diameter in the cytosolic and transmembrane regions, respectively (Fig. 1b). Monomer contacts are mediated by both the cytosolic and the transmembrane domains. The cytosolic domains are connected by i) a loose contact between the conserved TGES loop in the A domain (residues 231-235) and two α-helix termini in the P domain (Tyr541, Arg570 and Gln571) (Fig. 1a,b, Fig. S3a), and ii) a more extensive contact between the P domains of adjacent monomers, mediated via the auto-inhibitory R domain (Fig 1b, Table S2). The transmembrane interactions involve a number of functionally conserved hydrophobic residues in M3 of one monomer, and M7 and M10 of the adjacent monomer, augmented by two polar contacts from Ser316 and Asn317 in L3-4 to Gln780 (M7) and Arg859 (M10) (Fig. 1b, Fig. 2a, Table S2).

**Figure 2.**
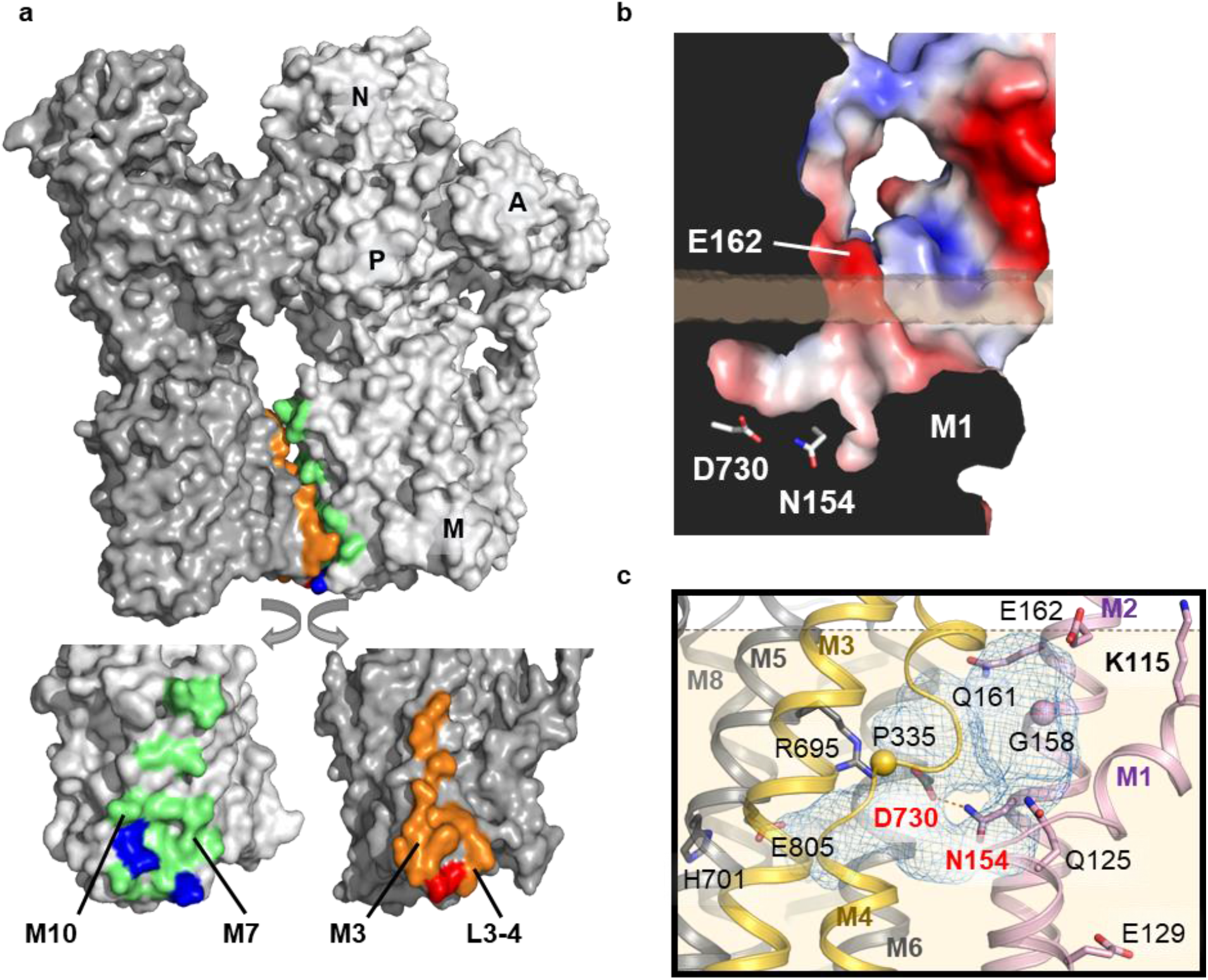
Structural features of Pma1. **a,** Interaction between M domains. Hydrophobic contacts are shown in orange (M3, L3-4) and light green (M7, M10), polar contacts in red and blue. **b,** A cross-section view of a Pma1 monomer perpendicular to the membrane plane, coloured by electrostatic surface potential (red, negative; blue, positive). A negatively charged proton entry funnel above M1 leads to the proton acceptor residue pair Asp730/Asn154 (shown as sticks). The funnel extends below the predicted lipid bilayer boundary45. **c,** Aqueous cavity between M1, M2, M6 and an unwound region of M4 (Pro335 shown as sphere). Important residues for proton transport and the snorkeling K115 (upper right corner) are shown as sticks (sphere for glycine). M1-2 are colored pink, M3-4 gold and M5-10 grey. In the enlarged view, the putative hydrogen bond (2.9 Å) between the proton acceptor/donor Asp730 and Asn154 (labelled in red) is shown as orange dashes. Arg695 is facing the cavity and could form a salt bridge with Asp730 (labelled in red) after proton release. The conserved residues His701 and Glu805 interact with each other in the *E*1 state.

The P-type ATPase conformational cycle is described by the *E*1/*E*2 scheme (Fig. S1b), with *E*1 denoting states with high affinity to the cytosolic substrate ion (H^+^ for Pma1). The structure is in an *E*1 conformation, resembling the AMPPCP-bound *E*1 structure of the plant proton pump AHA2^16^ and the pre-substrate binding, open-to-inside state of SERCA (Mg*E*1)^17^ (Fig. S3b). The conserved nucleotide binding pocket is occupied by MgADP (Fig. S3c). We have hence captured the basal state prior to ATP hydrolysis, occupied by excess MgADP.

Cytosolic access to the proton-binding site is open: the M1 helix with its characteristic 90° kink (induced by Pro123) is embedded deeply within the membrane, forming a wide funnel-shaped access towards the H^+^ binding site in M6, similar to the Ca^2+^ entry funnel in SERCA^17^ (Fig. 2b). The mouth of this funnel is negatively charged due to the conserved Glu162 in M2 (Fig. 2b), and it extends into a large aqueous cavity between M1, M2, M6, and an unwound region in M4 around Pro335 (Fig. 2c). The cavity is almost twice as large (~746 Å^3^) as in AHA2 (~380 Å^3^)^16^ and can accommodate up to 18 water molecules. The N-terminal part of M1 is strongly hydrophobic in Pma1, except for Lys115, which ‘snorkels’ toward the aqueous cytosolic phase (Fig. 2c). The C-terminal part of M1 contains two hydrophilic residues: Gln125 at the upper end, pointing into the protein interior, and Glu129 at the lower end, contacting the lipid environment. In line with its accessibility from the membrane centre, Glu129 is the site of dicyclohexyl carbodiimide (DCCD) binding and inhibition of Pma1^19^. DCCD might lock M1 in this open conformation, hence preventing proton occlusion and transport.

The proton-acceptor/donor in Pma1 is the conserved Asp730 in M6 (Fig. 2c). This residue corresponds to Asp684 in AHA2 and the Ca^2+^-coordinating residue Asp800 in SERCA and is essential for proton transport and *E*1→*E*2 transitions in Pma1^20,21^. Asp730 lies adjacent to the cavity and opposite the equally highly conserved Asn154 in M2, at a distance suitable for capturing a proton within an H-bond. Proton release to the extracellular side has been suggested to be driven by a conformational change that breaks this hydrogen bond and instead promotes the formation of a salt bridge between the aspartate and an adjacent arginine residue (Arg655 in AHA2), whilst concomitantly opening up the exit pathway between M1, M4, and M6^16,22^. The arginine has also been suggested to act as a positively charged ‘gate keeper’ preventing proton re-entry into the cavity. In Pma1, the equivalent of AHA2 Arg655 is His701 (M7), a fact that calls into question the similarity of the mechanisms of the two proton pumps. In our structure, His701 points away from the proton-binding site, interacting with Glu805 in M8 (Fig. 2c). The precise role of this residue pair is unclear, but both residues are necessary for proton pumping in Pma1^23^. The functional role postulated for Arg655 in AHA2 is likely taken over by Arg695 in Pma1, which is strictly conserved in fungi and positioned 2.5 helix turns up from His701, facing the ion-binding site (Fig. 2c).

The auto-inhibitory R domain extends from the end of M10 into a short region of disorder and then forms a mainly α-helical structure that lies adjacent to the P domain (Fig. 3a). It forms numerous interactions including a surprisingly large number of hydrophobic contacts, also involving the adjacent monomer’s P domain (Fig. 3b). The distance across the unmodelled gap between the M10 C terminus and the N termini of two adjacent R helices is very similar. The current assignment is based on a short extension of the R helix which points towards one of the two M10 termini (Fig. S3d).

**Figure 3.**
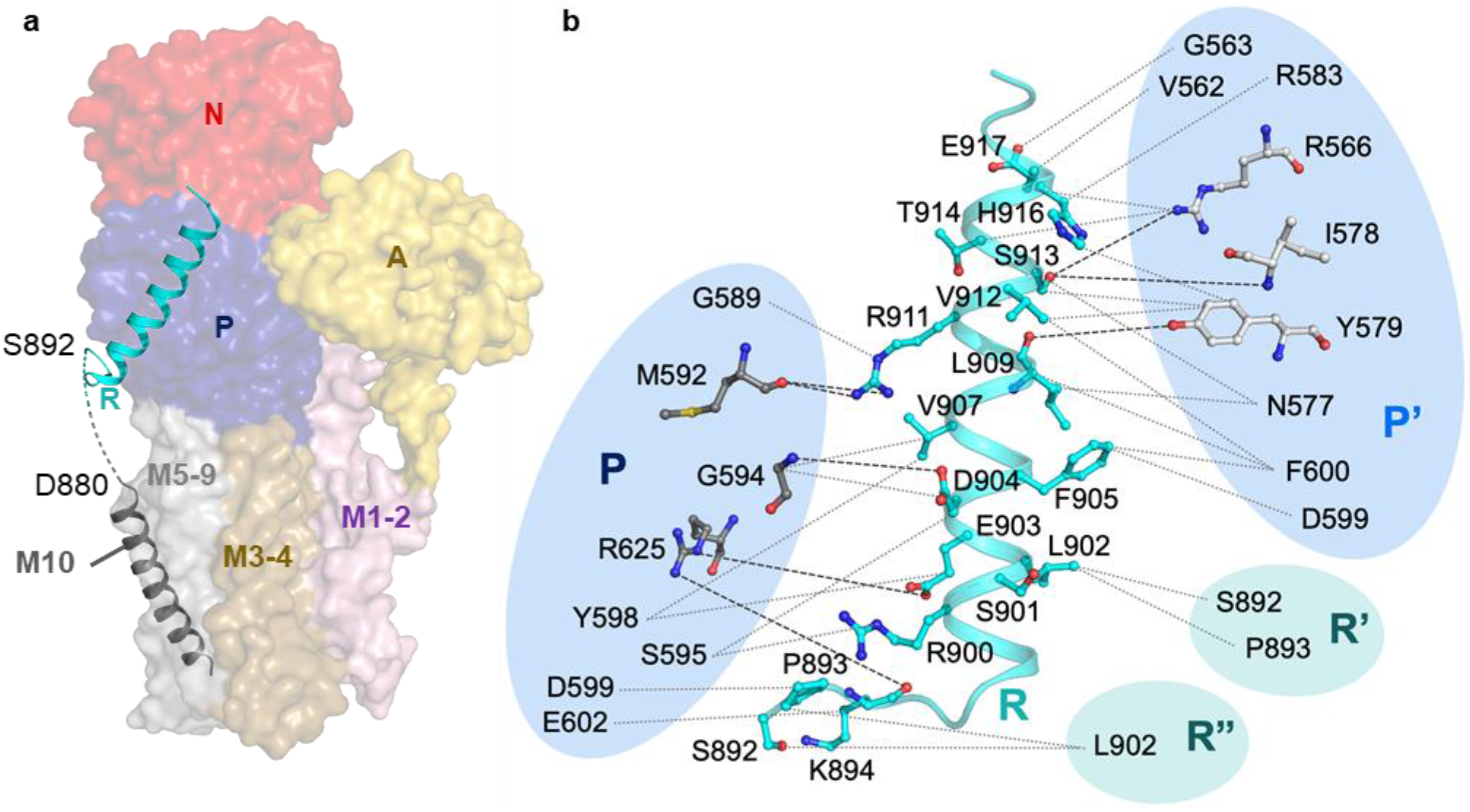
The autoinhibitory R domain and its interaction network. **a,** The R domain (cyan cartoon) is situated adjacent to the P domain (blue surface). It is connected to M10 (grey cartoon) via a brief region of disorder (dashed line). **b,** Schematic view of intra- and intermolecular interactions mediated by the R domain (cyan). Left side: intramolecular contact between the R and P domains, right side: intermolecular contact between the R domain and the adjacent monomer’s P domain (P’), and to two adjacent monomers’ R domains (R’, R”). P/P’ domain residues involved in polar contacts are shown as dark or light grey sticks, respectively.

To probe Pma1-lipid interactions in a native-like bilayer, we ran coarse-grained MD simulations of Pma1 in a complex membrane of different lipids occurring in *N. crassa*^13,24–26^ (Table S3). To prevent any bias of lipids situated inside the ring, one monomer was omitted, allowing free exchange during the simulations. Fractional interaction times of each lipid at the Pma1 surface suggest two putative binding sites at the monomer interface: Site I in the inner leaflet (involving M1, M3 and M10’) and Site II in the outer leaflet (between M4 and M10’), with a preferred binding of phosphatidylserine (PS) and phosphatidylcholine (PC), respectively (Fig. S4). Lipid-mediated hexamer contacts are in line with the observations that the hexamer is sensitive to treatment with certain detergents *in vitro* (Fig. S5), and that hexamer formation *in vivo* depends on an unperturbed lipid biosynthesis^11–13^. A local lipid density analysis shows that the accumulation of 16:2/18:2 PS (DIPS) at Site I is accompanied by an extended clustering of this lipid in the vicinity (Fig. 4a). 16:2/18:2 PC (DIPC) is the only other lipid that also accumulates at the interface, albeit to a lesser extent than DIPS, and with an otherwise even distribution around the protein (Fig. 4a, Fig. S6). Accumulation could either be driven by localised clustering of unsaturated tails, which would explain the presence of both DIPS and DIPC, or by clustering of the acidic PS headgroups, which has been observed previously^27^. Preferential binding of DIPS is in line with the dependency of detergent-purified Pma1 ATPase activity on acidic phospholipids^14^. Likewise, a recent study found PS enriched in SMA-solubilised Pma1^26^.

**Figure 4.**
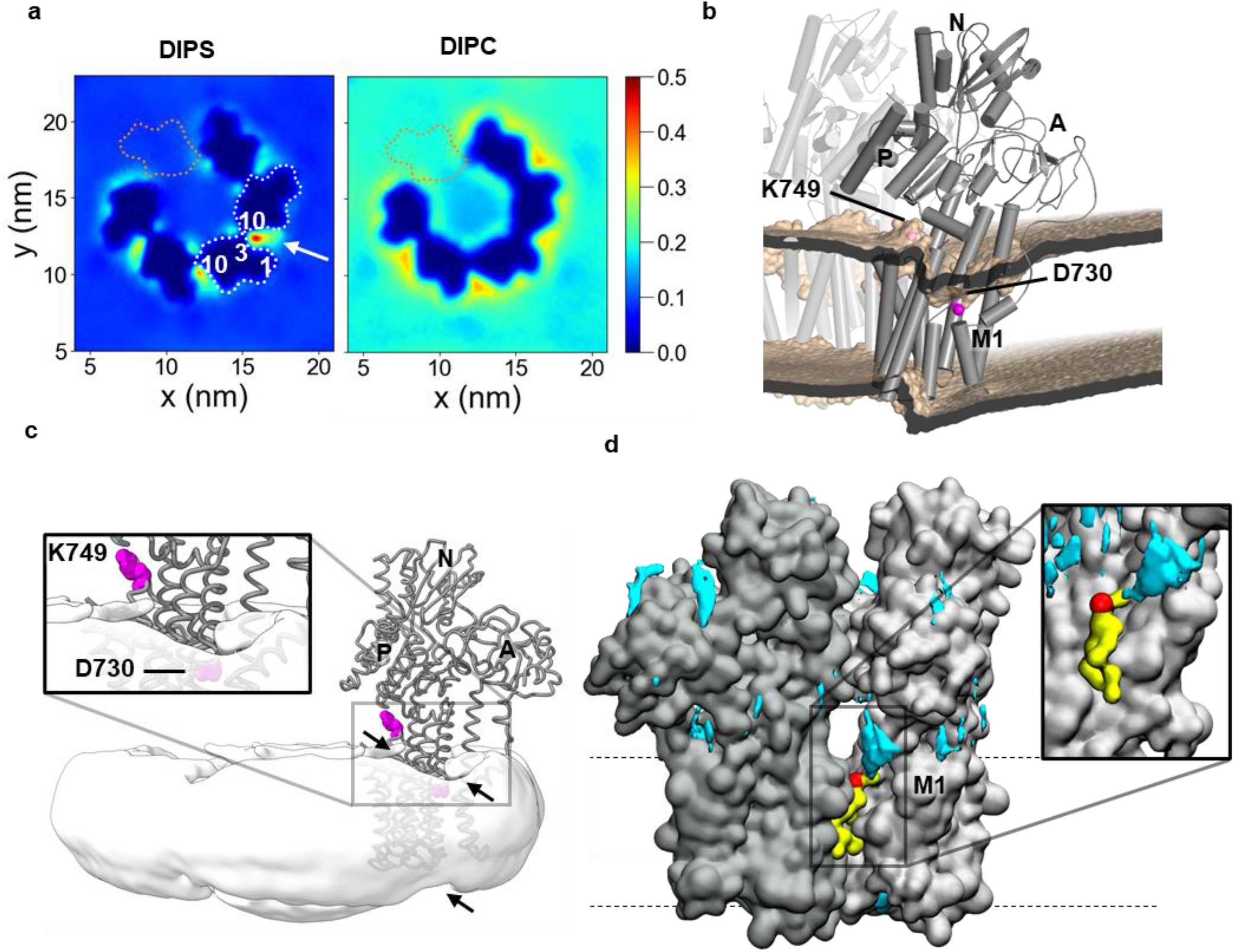
Lipid and cation interactions of Pma1. **a,** Simulated lipid densities for DIPS and DIPC. Arrow points at the pool clustering near the monomer interfaces. Values correspond to average number of molecules per nm3 and do not account for the respective membrane composition fraction. White dotted lines and numbers indicate monomer outlines and TM helix positions, respectively. Orange dotted lines indicate the omitted monomer. **b,** Side view of the deformation of the CG bilayer (wheat), shown as average phosphate position across 5 repeats. An atomistic Pma1 model was aligned for analysis. **c,** Cryo-EM map of hexameric Pma1, filtered to 12 Å, showing only the micelle after masking the protein map (Coot14). The Cα trace of one monomer is shown for reference, with the side chains of Asp730 and K749 highlighted in magenta. Arrows indicate local micelle distortion near the monomer interface. **d,** Locally increased density of Na+ ions (cyan) around the monomer interface. A representative DIPS molecule in Site I is shown in yellow, with phosphate group highlighted in red.

The simulations further suggest that the Pma1 protein has a considerable impact on the local membrane structure: near the M1-M4 bundle, the inner membrane leaflet forms a ‘dip’ that is most pronounced near the unwound region of M4 and along the outside-facing surfaces of M6 (containing the proton-accepting Asp730) and M9 (Fig. 4b, Fig. S7). Near this dip, on the inside of the ring, the membrane surface bulges upwards towards the P domain, probably attracted by the N-terminal end of M3 and by Lys749 in the L6-7 loop. These deformations are also evident from the cryo-EM map of the detergent/lipid micelle surrounding the Pma1 particles (Fig. 4c). This arrangement of negatively charged lipid head groups near the (also negatively charged, see Fig. 2b) mouth of the ion entry funnel could facilitate the attraction of cytosolic protons into the binding site. Indeed, the simulations show an increased local density of Na^+^ ions in this region. (Fig. 4d). The simulations suggest a ~10Å decrease in membrane thickness (from ~37Å to ~27Å) near the proton entry funnel and a local thickening to ~54Å in the membrane region below the P domain (Fig. 5b, Fig. S10). Mutual adaptations between protein and membrane, as well as membrane deformations assisting cytosolic ion entry have already been suggested for SERCA^17,29^ and might be a common feature of P-type ATPase function.

To develop a model for the catalytic cycle of Pma1, we generated a series of SERCA-based homology models. The transition state of phosphorylation, *E*1~P, is linked to ion occlusion within the M domain^30,31^. Interestingly, in this state Asp730 moves away from Asn154, implying a break of the suggested hydrogen bond (Fig. 2c). Nevertheless, the predicted pK_a_ value of Asp730 in this state (7.2)^32,33^ suggests that it remains protonated, likely because it is buried inside the hydrophobic core of the protein. The backbone carbonyl oxygens of Ile332 and Ala729, may act as stabilising H-bond acceptors. Arg695, the conserved residue suggested to be involved in proton repulsion in *E*2, takes a position that shields Asp730 from the entry channel and possibly prevents the acquired proton from escaping back to the cytosol.

In the open-to-outside *E*2P model, Asp730 lies at the inner end of the exit funnel, which is lined by predominantly hydrophobic side chains, except for Asp143 approximately halfway down the funnel. Both His701 and Arg695 are near Asp730, and the Arg695 side chain can freely rotate into bonding distance (Fig S6a). It could hence prevent re-protonation of Asp730 before the exit pathway closes, acting as an in-built counter-ion to neutralise Asp730 during the *E*2→*E*1 transition, as suggested for AHA2^16,22^. A conspicuous cluster of acidic residues Glu139, Asp140 (L1-2), Asp143 (M2), Glu324 (M4) and Glu720 (M6) lies at the extracellular end of the funnel (Fig. S6a), likely facilitating proton extrusion.

In our structure, the R-helix interacts exclusively with the P domain which is expected to move relatively little during proton pumping. All modelled conformations can accommodate the R-helix in this position, in line with the observation that binding of free R-peptide permits Pma1 activity^10^. However, the distance between the C terminus of M10 and the N terminus of the R helix increases from 30 Å in the *E*1 structure to 41 Å in the *E*2 homology model (Fig. S6b), supporting the ‘clamping’ model of auto-inhibition^5^, as the 16-residue linker between M10 and the R-helix might restrain the *E*1-*E*2 transition.

In yeast, release of auto-inhibition upon glucose-sensing has been linked to phosphorylation of Ser899 (Ser901 in *N. crassa*) by Ptk2^34^ and tandem phosphorylation of Ser911/Thr912 (Ser913/Thr914 in *N. crassa*) by an unknown kinase^35^. Strikingly, neither of these residues is located at the binding interface of the *cis*-acting R domain (Fig. 3b). A release of the R domain from the P domain in a monomeric context would therefore need to involve phosphorylation-induced structural alterations within the R-helix itself, causing disruption of R-P interactions. Such a mechanism seems rather unlikely and points towards a physiological role of the Pma1 hexamer. While monomeric Pma1 is capable of pumping protons^38^, activity *in vitro* is mostly reported in a hexameric context, and the physiological importance of multimeric plasma membrane H^+^-ATPases is becoming increasingly evident^39,40^. Studies with purified fungal membranes have indicated the presence of either two cooperative ATP-binding sites or a larger number of sites with a lower degree of cooperativity^41^. Inhibition studies suggest that binding of one DCCD molecule blocks ATP hydrolysis in ~2.5 molecules of Pma1^19^, suggesting a physiological oligomer. Hexameric assemblies of the homology models show that there is sufficient space for the expected cytosolic domain movements throughout the catalytic cycle (Fig. S6c). None of the 20 possible neighbouring conformations clash (assuming an unbound R domain, see below), providing no support for any cooperativity between neighbouring monomers. The models further suggest that the hexameric ring would need to expand slightly to accommodate the opening of the extracellular pathway, or alternatively that Pma1 does not open up as widely as SERCA. This may reflect the smaller size of the transported ion and the need to ensure quick closure upon proton release to prevent back-flow at high membrane potentials.

The domain packing of the auto-inhibited state is tighter than in any of the homology models, reflected in the smallest accessible surface area and smallest outer diameter compared to the models (Fig. S6c,d). The R domains are sandwiched between adjacent P domains, forming an extensive interaction network (Fig. 1a, Fig. 3b). R domains of adjacent monomers interact directly, thereby linking all R domains within the hexamer. The R domain-mediated cross-links could inhibit P-domain separation during pumping and hence augment the auto-inhibitory effect. Furthermore, in four of the possible neighboring combinations within the hexamer (*E*2:*E*1, *E*2:*E*1~P, *E*2P:*E*1 and *E*2~P:*E*1), the R helix cannot be accommodated in the position observed in the *E*1 state. This suggests that the R domain gets displaced from the P domain upon Pma1 activation.

Phosphorylation of Ser913/Thr914 would disturb the *trans*-contact between the R domain and the adjacent P domain. In contrast, Ser901 points into the centre of the ring, making its phosphorylation unlikely to affect the intermolecular contact. A single molecule of Ptk2 can fit into the centre of the Pma1 hexamer, and could initially phosphorylate Ser901 in one monomer, followed by ATP binding and hydrolysis. The accompanying domain rearrangements would then increase accessibility of the neighboring monomers’ R domains, thus perpetuating the activation through the hexamer. Notably, the Ser901 sidechain is situated only 5.4 Å away from Arg900 of one adjacent monomer, allowing for an R-R contact upon phosphorylation which could eventually sequester together all six released R domains. This could aid the activation process and stabilise the activated state. A similar interaction between inhibitory domains was recently suggested for AHA2^40^.

To explore the amenability of Pma1 to inhibition by small-molecule drugs, we mapped the conservation level of the *N. crassa* Pma1 sequence relative to the most important human- and plant-pathogenic fungi (see *Methods* for species) on the protein surface^42^ (Fig. S7a). Focusing on the M domain, which would be most accessible, we identified a deep groove between M1, M3 and M4, along a ridge of highly conserved residues, and extending to the extracellular surface (Fig. S7a). To probe this groove’s suitability for drug binding, we docked a set of nine previously reported Pma1 inhibitors^43^ into the structure, using the entire M domain of one monomer as search space. Of the 162 calculated docking poses, 28 had a calculated binding affinity of < −8.5 kcal/mol and were not located in the hexamer interface. Encouragingly, 18 of these 28 docking poses occupied the conserved groove described above (Fig. S7b). Structures of Pma1-inhibitor complexes will be needed to confirm the binding experimentally. Notably, most of the conserved residues around this groove are also conserved in the proton pumps of many economically important plants, making a targeting of this site potentially more promising for therapeutic applications than for plant protection.

In conclusion, we identified core elements of known proton-pumps in Pma1: a central proton acceptor/donor (Asp730), a positively charged residue to control pK_a_ changes of the proton acceptor/donor (Arg695), and bound water to facilitate proton transport^44^. The hexamer explains the dependency of Pma1 activity on anionic lipids by a preferential interaction site at the monomer interface. Anionic lipids could also be involved in the attraction of protons into the acceptor site, as suggested by a local membrane deformation and clustering of cations. We find no structural basis for cooperativity within the hexamer, but the structure implies that auto-inhibition is enhanced – if not even dependent on – a hexamer context. Activation by phosphorylation likely occurs sequentially, and could be accelerated by mutual sequestering of the released R-domains. The Pma1 structure is a major advance towards the development of specific inhibitors.

## Material and Methods

### Plasma Membrane Preparation

Purification of Pma1 was carried out from native source, using a cell wall-less *Neurospora crassa* strain (FGSC 4761). Cells were grown for 24 h at 30°C and 185 rpm (Brunswick Innova 44) in Vogel’s medium supplemented with 2% (w/v) mannitol, 0.75% (w/v) yeast extract (Difco), and 0.75% (w/v) nutrient broth (Difco) using 2.5 L Ultra Yield™ flasks with AirOtop™ lids. Plasma membrane preparation was carried out as described previously^46^ with some adjustments to the original protocol. All centrifugation steps were conducted at 4°C, cells and membranes were kept on ice unless stated otherwise. The cells were harvested by centrifugation (15 min at 700˟*g*) and washed four times with ice-cold Buffer A (50 mM Tris pH 7.5, 10 mM MgSO_4_, 250 mM mannitol) (15 min, 700˟g) before they were agglutinated with 0.5 mg/mL Concanavalin A in Buffer A for 10 min at room temperature (RT) and another 10 min on ice. Agglutinated cells were pelleted for 6 min at 200˟g and washed once with ice-cold buffer A. Lysis was obtained by homogenizing the cells in ice-cold lysis buffer (10 mM Tris pH 7.5, 1 mM MgCl_2_, 1 mM CaCl_2_, 1.5 μg/mL chymostatin, 1 μg/mL DNase I) containing 1.2% sodium deoxycholate (DOC) using a 50 cm^3^ glass homogenizer. After centrifugation (30 min, 14000˟g), the membrane pellet was first washed with Buffer B (10 mM Tris pH 7.5, 1.5 μg/mL chymostatin) containing 0.6% DOC and then with DOC-free Buffer B. Dissociation of Concanavalin A from the plasma membrane was achieved by incubation with 0.5 M α-methylmannoside in Buffer B for 5 min at 30°C. The dissociated membranes were diluted with ice-cold Buffer B, pelleted (30 min, 14000˟g) and washed once with ice-cold Buffer B. After resuspension in ice-cold Buffer B, membranes were flash frozen in liquid nitrogen and stored at −80°C until protein purification.

### Detergent Solubilization & Protein Purification

Membranes were thawed in a RT water bath, spun down (30 min, 17700˟g, 4°C) and washed twice (20 mM Hepes pH 7.2, 0.5 M NaCl, 1 mM PMSF). Solubilization was carried out for 1 h at 12 to 19°C with 2.25 mg/mL of the detergent β-dodecyl-maltopyranoside (DDM) in Pma1 solubilisation buffer (10 mM Tris pH 7.5, 150 mM NaCl, 1 mM Na_2_-ATP, 5 mM MgSO_4_, 0.1 mM Na_3_VO_4_, 2 μg/mL chymostatin) using a total membrane protein concentration of 2 mg/mL determined by BCA assay. All further purification and centrifugation steps were performed at 4°C. Unsolubilized membranes were removed by centrifugation (50 min, 17700˟g) and the supernatant was filtered through a 0.45 μm nylon membrane.

The protein solution was concentrated by pressure dialysis with a stirred cell (Amicon) and a 300 kDa MWCO filter membrane, diluted to a NaCl concentration of ~40-45 mM (50 mM MES/Tris pH 7, 5 mM MgSO_4_, 200 g/L glycerol, 1 mM DTT, 2 mM EDTA), and loaded onto a Q-HP anion exchange (AEX) column (GE Healthcare) equilibrated with dilution buffer containing 0.1 mg/mL DDM. The AEX column was washed with 20 column volumes (CV) wash buffer (50 mM MES/Tris pH 7, 20 mM KCl, 5 mM MgSO_4_, 1 mM DTT, 2 mM EDTA, 0.15 mg/mL DDM, 2 μg/mL chymostatin) and eluted with a linear gradient to 500 mM KCl over 20 CV. Peak elution fractions were pooled, diluted 1:10 (50 mM MES/Tris pH 6, 5 mM MgSO_4_, 2 mM EDTA, 1 mM DTT, 2 μg/mL chymostatin, 0.15 mg/mL DDM) and concentrated to a volume of 1-2 mL with Amicon Ultra centrifugal filters (100 kDa MWCO).

The concentrated sample was loaded onto a glycerol density gradient from 20-40% (w/v) (5% steps) and ultra-centrifuged for 16 h at 34,400 g (Optima L-100 XP, Ti70 rotor, acceleration 2, no brake). The gradient was harvested in 1 mL fractions from bottom to top using a peristaltic pump with HPLC tubing (i.d. 0.15 mm). The purest fractions were pooled, diluted 1:10 in glycerol-free buffer and concentrated to a volume of 0.5-1 mL. The remaining Pma1-containing fractions were also pooled, diluted 1:5 and concentrated to a volume of 0.5-1 mL. The concentrated samples were loaded onto two separate glycerol density gradients from 20-45% (w/v) (5% steps), and ultra-centrifuged and harvested as described above. The purest fractions of both gradients were pooled, concentrated and loaded onto a Superose 6 Increase 10/300 (GE Healthcare) size-exclusion chromatography (SEC) column equilibrated with SEC buffer (50 mM MES/Tris pH 6.5, 200 g/L glycerol, 50 mM KCl, 5 mM MgSO_4_, 2 mM EDTA, 1 mM DTT, 2 μg/mL chymostatin, 0.15 mg/mL DDM) Peak fractions were evaluated by silver stain SDS-PAGE and negative stain electron microscopy.

### Negative Stain EM

Samples were diluted to 0.015 mg/mL (determined with a NanoDrop™ One and Abs 0.1% = 1.044 calculated by ProtParam^47^) with glycerol-free SEC buffer. Carbon-coated 400-mesh copper grids were glow-discharged (15 mA, 45 s), incubated with 3 μL sample for 30 s and stained twice for 15 s with 3 μL 2% uranyl formate. Blotting after each incubation step was performed at a 45° angle with Whatman filter paper (#1). The grids were imaged with a 120 kV Tecnai™ Spirit (Thermo Fisher) at 52’000 x magnification.

### Cryo-EM

Suitable SEC elution fractions (see Fig. S4e) were pooled and concentrated to 40 μL at 3 mg/mL with Amicon Ultra centrifugal filters (100 kDa MWCO). The buffer was exchanged to cryo-EM buffer (30 mM MES pH 6.5, 1 mM MgSO_4_, 1 mM DTT, 2 μg/mL chymostatin, 0.15 mg/mL DDM) with a 0.5 mL Zeba Spin column (7 kDa) following the manufacturer’s protocol. The eluted sample was concentrated to 3.7 mg/mL and incubated for 2 h on ice with 1 mM ADP and 0.6 mM AlF_4_ at a final protein concentration of 2 mg/mL (~20 μM). Cryo-EM grids were prepared by applying 3 μL sample to twice glow-discharged (15mA, 45s) C-flat R2/2 400-mesh copper grids which were blotted for 11 s (10°C, 70% humidity, S&S 595 filter paper) and plunge-frozen in liquid ethane using a Vitrobot Mark IV (Thermo Fisher). Cryo-EM grids were stored in liquid nitrogen until data were collected with a 300 kV Titan Krios G3i (Thermo Fisher) equipped with a K3 direct electron detector (Gatan). 2780 micrographs were recorded automatically in EPU (Thermo Fisher) in counting mode at 105’000 x magnification with a calibrated pixel size of 0.837 Å. The defocus range was set from −1.3 to −2.5 μm and each micrograph was dose-fractionated to 50 frames with a total exposure time of 3 s, resulting in a total dose of ~42 e^−^/Å^2^ (see Table S1).

### Cryo-EM Image Analysis

CryoSPARC^48^ was used for particle picking and selection, while all further refinement steps were performed in Relion^49–51^. After patch-based motion correction and contrast transfer function (CTF) estimation in cryoSPARC, particles were picked with the blob picker (180 Å diameter) and manually curated, resulting in 483’742 particles. Four iterative rounds of 2D classification led to 95’525 particles, which were used for an *ab initio* reconstruction with three classes, resulting in two protein classes with 81’009 particles and one junk class. A homogenous refinement gave an initial map of hexameric Pma1 at 3.6 Å resolution (using C1 symmetry). The raw micrographs were then imported to Relion 3.0 and motion corrected with MotionCor2^52^. CTF estimation was performed using Gctf^53^ on non-dose weighted aligned micrographs and the cryoSPARC particles were re-extracted using the coordinates from the homogenous refinement. The following processing steps were performed in Relion 3.1-beta. An initial 3D refinement applying C6 symmetry resulted in a 3.79 Å map. CTF refinement and anisotropic magnification estimation^54^ improved the resolution to 3.53 Å, Bayesian polishing^55^ to 3.42 Å (map A) and another CTF refinement to 3.28 Å (map B). The map was of high quality for the M and P domains and part of the R domain, but the A and N domains were less well resolved.

To improve the resolution of the cytosolic A and N domains, the particles of maps A and B were symmetry-expanded and a 3D classification focused on the monomer^50^ was performed with each set. The best monomer classes comprised 43.8% (214’647) and 59.9% (293’999) of the expanded particles, respectively, and were used for 3D refinement (applying C1 symmetry and only local angular searches^50^). The refinement was started with a mask encompassing the whole hexamer and continued from a late iteration using a mask that covered only one monomer. Focused refinement improved the map resolution to 3.25 Å (map a) and 3.21 Å (map b), respectively, and significantly improved the density for the cytosolic A and N domains, allowing for model building.

Due to the high number of discarded particles after the monomer-focused 3D classification, the (not expanded) hexamer particles from map B were subjected to a 3D classification focused on the rigid part of the hexamer (M and P domains). The remaining 59’511 particles (73.3%) were 3D refined (with a mask around the whole hexamer) to 3.28 Å resolution (map C). The map was postprocessed with DeepEMhancer for sharpening^56^.

To further improve the monomer map, the particles of map C were symmetry-expanded and another monomer-focused 3D classification was performed using a mask without the flexible A domain. The best monomer class comprised 71.9% (261’196) of the expanded particles and was 3D refined as described above to a resolution of 3.18 Å (map c). The three monomer maps a, b, and c were combined using phenix.combine_focused_maps^57^; rigid-body fitting of the model was disabled. The same was done for the corresponding half maps, which were subsequently density-modified with phenix.resolve_cryo_em^58^. Four different settings were applied in the density modification: sharpened or unsharpened half maps were used, in both cases with the default FSC-based final resolution-dependent scaling enabled or disabled. The four density-modified composite maps were then combined into the final monomer map (map d) with a resolution of 3.11 Å, estimated with the *Phenix Comprehensive validation* tool^57^. A summary of the refinement process is depicted in Fig. S2.

### Model Building and Refinement

Monomer maps a, b and c were used for *de novo* model building in Coot^59^, assisted by homology models (see below) and the Namdinator server^60^. Amino acid residue assignment was based on clearly defined densities of bulky residues (Phe, Trp, Tyr and Arg), the MgADP ligand and K^+^ ion were added to the model based on the crystal structures of the plant proton pump^16^ and SERCA^17, 30^. The final model was refined using phenix.real_space_refine^61^ and the density-modified monomer map d. C6 symmetry was applied to expand the monomer model into the density-modified hexamer map C using UCSF Chimera^62^. The geometry of both models was validated using MolProbity^63^ implemented in the *Phenix Comprehensive validation* tool (see Table S1). Final models of the monomer and hexamer were submitted to the Protein Data Bank, PDB ID 7NXF and 7NY1, respectively.

### Multiple Sequence Alignments

*Neurospora crassa* Pma1 (accession code sp|P07038) was aligned using ClustalOmega to the plasma membrane proton pumps of the following organisms: i) human pathogenic fungi: *Sporothrix schenckii* (tr|A0A0F2M765), *Histoplasma capsulatum* (sp|Q07421), *Coccidioides immites* (tr|A0A0E1RVX1), *Blastomyces dermatitidis* (tr|T5BWH0), *Acremonium chrysogenum* (tr|A0A4Y6GPG7), *Trichophyton rubrum* (tr|A0A022W272), *Candida glabrata* (tr|Q6FXU5), *Candida auris* (tr|A0A2H0ZKV9), *Candida albicans* (sp|P28877), *Pneumocystis jirovecii* (tr|A0A0W4ZIS2), *Aspergillus fumigatus* (tr|Q96TH7), *Talaromyces marneffei* (tr|A0A093Y2U3), *Syncephalastrum racemosum* (ORZ00574.1), *Rhizopus stolonifera* (RCI02493.1), *Lichtheimia corymbifera* (CDH55078.1), *Cryptococcus gattii* (tr|A0A0D0U934), *Cryptococcus neoformans* (tr|O74242); ii) plant pathogenic fungi: *Claviceps purpurea* (CCE32805.1), *Colletotrichum gloeosphorioides* (KAF3800644.1), *Magnaporthe oryzae* (ELQ65709.1), *Fusarium oxysporum* (RKL39108.1), *Botrytis cinerea* (EMR90163.1), *Fusarium graminearum* (PCD22889.1), Aspergillus niger (GAQ33769.1), *Blumeria graminis* (AAK94188.1), *Sclerotinia slerotiorum* (tr|A7F838), *Mycosphaerella graminicola* (XP_003852209.1), *Cochliobolus heterostrophus* (tr|M2TMB0), *Rhizoctonia solani* (tr|A0A0B7FM75), *Ustilago maydis* (XP_011388984.1), *Puccinia graminis* (tr|A0A5B0P5Y0); iii) plant proton pumps: *Coffea eugenioides* (XP_027176212.1), *Spinacia oleracea* (XP_021865157.1), *Cucumis sativus* (XP_004152192.1), *Hordeum vulgare* (KAE8805265.1:29-858), *Jatropha curcas* (XP_012068768.1:37-846), *Triticum aestivum* (P83970.1:29-858), *Ananas comosus* (XP_020090190.1), *Chenopodium quinoa* (XP_021755229.1:32-855), *Carica papaya* (XP_021899224.1), *Ricinus communis* (XP_015572514.1:34-875), *Punica granatum* (XP_031378860.1), *Nicotiana tabacum* (NP_001312285.1), *Brassica napus* (XP_022556197.1), *Gossypium australe* (KAA3489374.1:33-874), *Arabidopsis thaliana* (NP_194748.1), *Manihot esculenta* (XP_021598156.1:34-875), *Malus domestica* (XP_008372282.1:35-876), *Camellia sinensis* (XP_028098451.1:32-870), *Zea mays* (AQK46772.1:25-866), *Theobroma cacao* (EOY29625.1), *Sesamum indicum* (XP_011084025.1), *Hevea brasiliensis* (XP_021654241.1:37-846), *Glycine max* (XP_003549696.1:32-903)

### Homology modelling & Internal Cavities

Homology models were generated with SWISS-MODEL^64^, based on a structure-based alignment^65^ and the SERCA crystal structures 1T5T (*E*1~P)^30^, 3B9B (*E*2P)^66^, 3N5K (*E*2~P)^67^, and 3NAL (*E*2)^68^. The SWISS-MODEL homology models were aligned to monomeric and hexameric Pma1 via M6-M10. Internal cavities of monomeric Pma1 and homology models were detected with HOLLOW^69^.

### Inhibitor Docking

The docking of a set of tetrahydrocarbazole (THCA) Pma1 inhibitors^43^ into the structure of the Pma1 monomer was done with AutoDock Vina^70^. Nine THCA compounds were added in both their R and S conformations to a box emcompassing the entire transmembrane domain of Pma1. Nine binding modes per conformation were analysed, resulting in a total of 162 binding events of which 60 displayed affinities stronger than −8.5 kcal/mol. 39 binding events occurred at one of the monomer interfaces and were therefore excluded from further analysis. 3 binding events took place within the proton entry funnel, 18 within the exit funnel. The strongest binding, exceeding −9 kcal/mol, was observed for 10 events within the exit funnel. Residues involved in the putative binding of the respective THCA compounds (9S, 10S/R, 11R) were identified using LigPlot+^71^ (see Fig. S7b).

### Coarse-grained Molecular Dynamics Simulations

The pentamer structure coordinates were converted to coarse-grained Martini representation using the martinize script^72^. Their tertiary structures were constrained using the ElNeDyn elastic network between all chains^73^ with a force constant of 500 kJ/mol/nm^2^ and a cut off of 0.9 nm. The CG protein coordinates were then positioned in the centre of a simulation box of size 23 x 23 x 23 nm^3^ with its principal transmembrane axis aligned parallel to the z axis and embedded in a complex asymmetric membrane bilayer comprised of 14 lipid species using the insane script^74^. The membrane bilayer was built according to Table S3. To investigate the role of lipid saturation, we also set up simulations of pentameric Pma1 in a 50% PIPC: 50% DIPC symmetric membrane. 0.15 M NaCl was added to the solvated system. The Martini coarse-grained force field version 2.2^72^ was used for protein and version 2.0 for lipids. All the simulations were performed using Gromacs 2019.1^75^. The systems were energy minimised using the steepest descents method, equilibrated for 50 ns using 20 fs timesteps in the isothermal-isobaric (NPT) ensemble at 323 K using the V-rescale thermostat^76^ and at 1 bar using a semi-isotropic Berendsen barostat^77^. Five production simulations were run to 10 μs using the Parrinello-Rahman barostat^78^. Data were analysed using VMD^79^, Gromacs tools^77^ and in-house scripts. A weighted atomic density for Na^+^ at each gridpoint was calculated using the VMD Volmap tool over 50 μs of simulation with default settings. Plots were made using MDAnalysis^80^ and Microsoft Excel.

## Figure preparation

Figures were prepared using VMD^79^, ChimeraX^81^ and PyMOL^82^. The OPM database^45^ was used to estimate the membrane position for Fig. 2b.

## Data and materials availability

Coordinates and EM maps have been deposited in the Protein Data Bank and EM Data Bank with accession numbers as follows: Monomer, 7NXF and EMD-12638/-12641/-12642/-12643; hexamer, 7NY1 and EMD-12644. Correspondence and requests for materials should be addressed to M.B.

## Acknowledgments

We thank Poul Nissen, Nikolaj Düring Drachmann, Dorota Focht, Christina Grønberg and Marco Mazzorana for contributions to initial project design and purification strategy, and Richard Henderson for help with establishing suitable cryo-EM conditions. We are grateful to Susann Kaltwasser, Simone Prinz, Sonja Welsch and Julius Demmer for technical assistance and help with data collection and analysis. We thank Janet Vonck for help evaluating preliminary cryo-EM data collected for the project, and Gerhard Hummer for discussions on simulation data analysis. Funding was provided by a Springboard Award from the Academy of Medical Sciences, the John Fell Fund 152/059, the E P A Cephalosporin Fund CF346, the Wellcome Trust Graduate programme in Cellular Structural Biology ref. 220063/Z/20/Z, a FEBS Short-Term Fellowship, the ERASMUS+ Staff Training Mobility programme, the Biochemical Society and the Max-Planck Society.

## Author contributions

S.H. performed Pma1 purification, collected and processed the data, and determined, refined and analysed the structures. M.M.G.G. conducted and analysed MD simulations. B.J.M. assisted with data processing, R.A.C. with MD simulations, and D.J.M. with data collection. W. K. provided cryo-EM resources and co-supervised the project. M.B. conceptualised and supervised the project, analysed the structure, and wrote the paper. All authors reviewed and edited the manuscript.

## Additional information

Correspondence and requests for materials should be addressed to.

