## Supplementary Information for "Structure of the hexameric fungal plasma membrane proton pump in its auto-inhibited state"

**Affiliations:**

[3]Deryck Mills passed away on 7 July 2020.

**Supplementary Figures:**

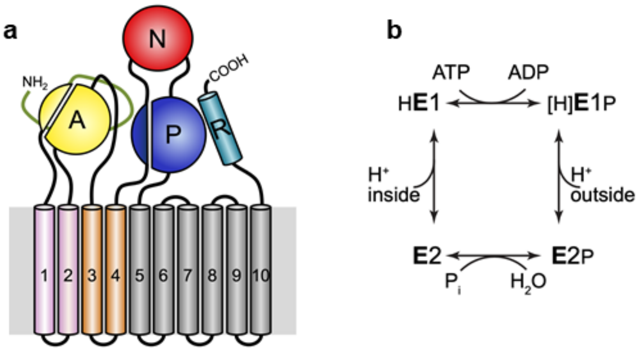

**Supplementary Figure S1 | Topology diagram and E1/E2 scheme of Pma1.**

**a,** Overall Pma1 topology. Nucleotide-binding (N) domain red, actuator (A) domain yellow, phosphorylation (P) domain blue, regulatory (R) domain cyan, N-terminal extension (green), M1-2 pink, M3-4 gold, and M6-10 grey. **b,** Canonical *E*1-*E*2 catalytic cycle for proton pumping by Pma1 with transient phosphorylation.

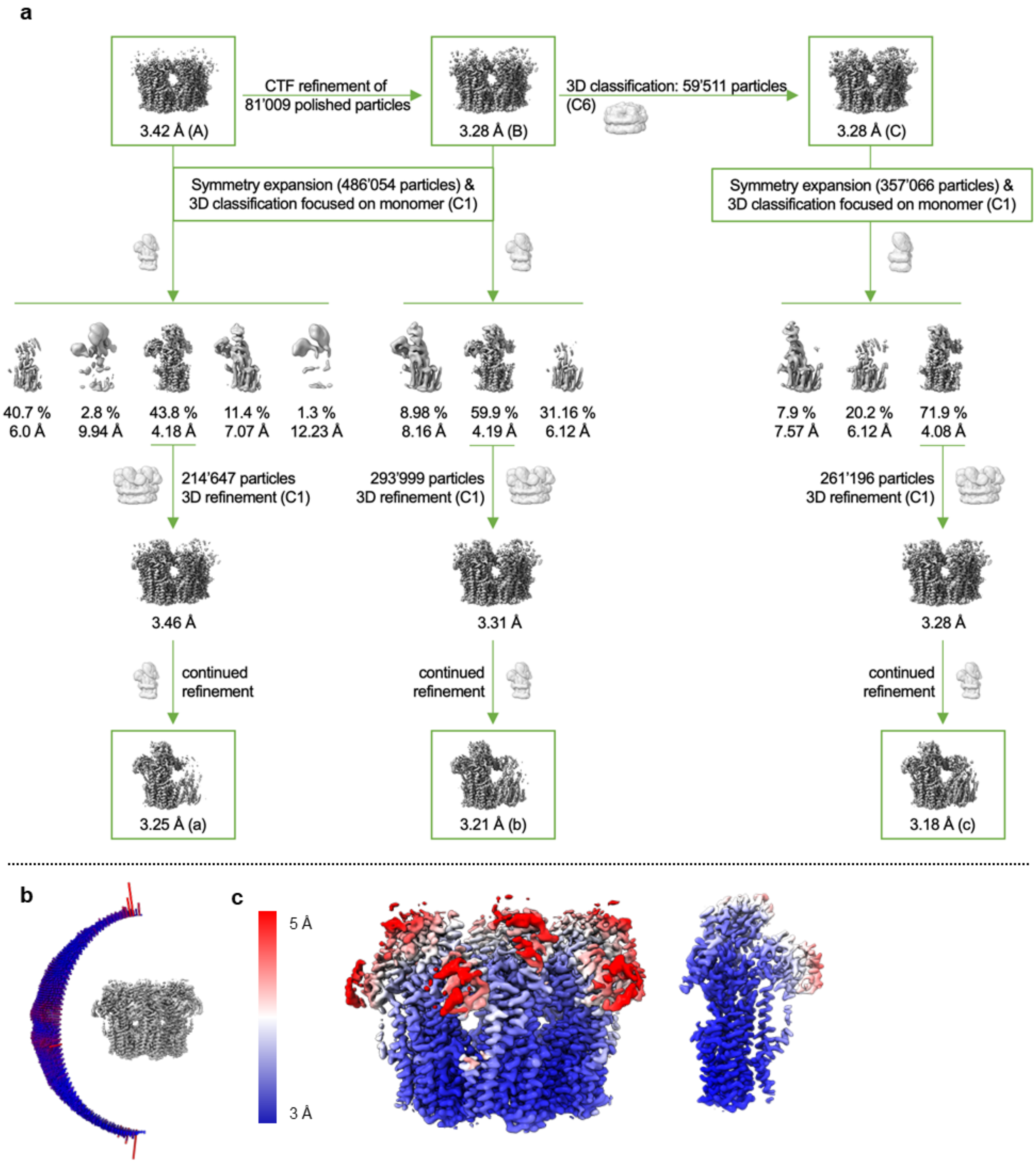

**Supplementary Figure S2 | Cryo-EM data processing and map attributes.**

**a,** Refinement workflow resulting in the final hexamer map (C) and the focused monomer maps (a), (b) and (c). **b,** Angular distribution plot of particles that contributed to the final hexamer map. The height of the bars is proportional to the number of particles in those views. **c,** Local resolution of the final hexamer map (C) and the combined monomer map (d). Resolution estimates from Relion 3DRefine[6].

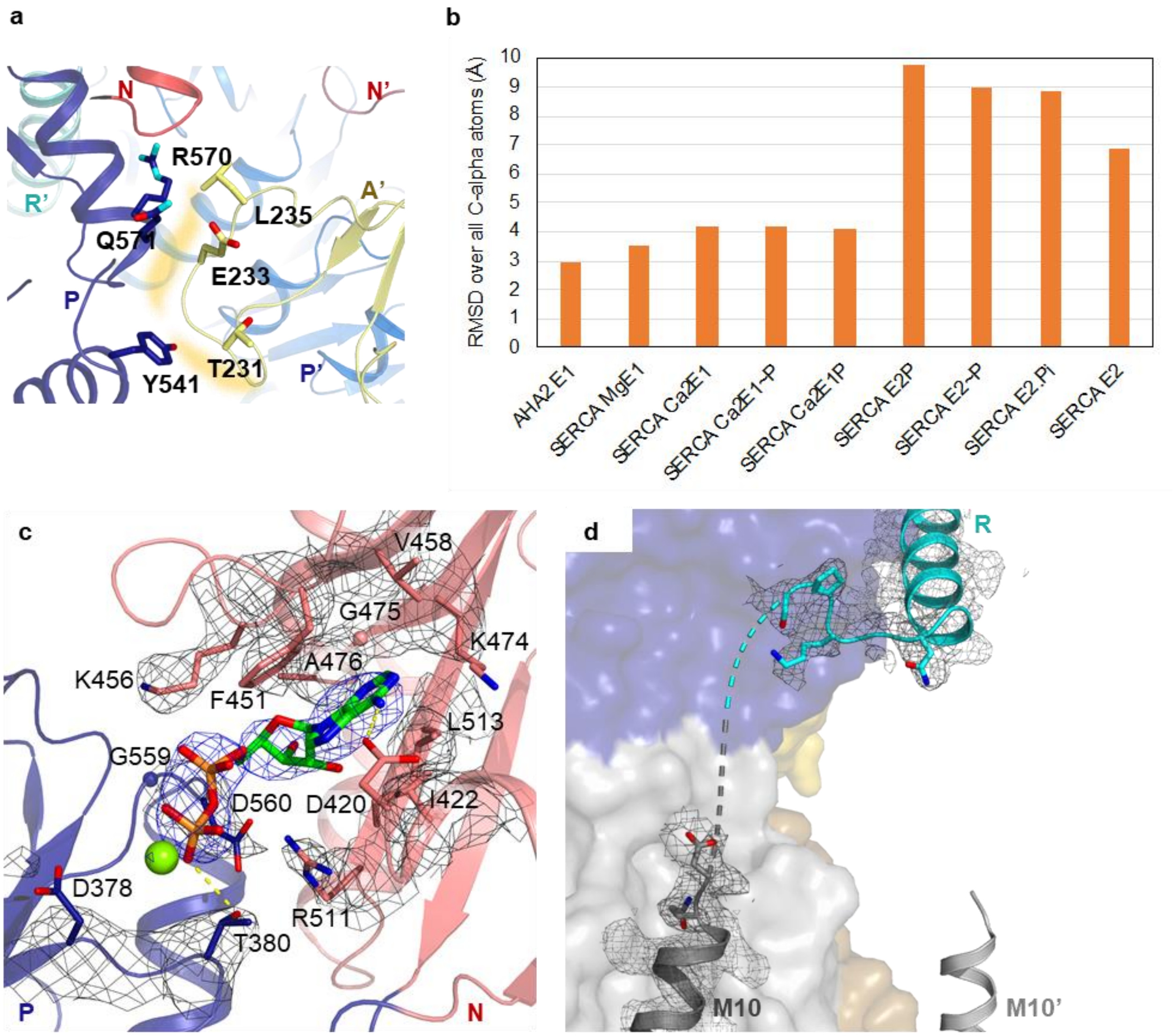

**Supplementary Figure S3 | Structural state comparison of Pma1 with related P-type ATPases.**

**a,** Interaction interface between the P domain (blue) and the TGES loop of the adjacent monomer’s A domain (A’, yellow). Contact area is shown as yellow shadow, involved side chains are shown as sticks.

**b,** RMSD (root mean square deviation) calculated over all C-alpha atoms of Pma1 compared to the crystal structures of AHA2 (*E*1) and SERCA in different states along the catalytic cycle using PyMOL[42]. PDB entries: 5KSD (AHA2 E1), 4HW1 (SERCA Mg*E*1), 3N8G (SERCA Ca2*E*1), 1T5T (SERCA Ca2*E*1~P), 3BA6 (SERCA Ca2*E*1P), 3B9B (SERCA *E*2P), 3N5K (SERCA *E*2~P), 1WPJ (SERCA $E2.P_i$), 3NAL (SERCA *E*2). **c,** ADP is bound at the nucleotide-binding site between the P (blue) and N (light red) domains. Residues involved in ADP coordination and

Asp378 are shown as sticks (spheres for glycine) with C-atoms coloured in green (ADP) or according to their domain, a $Mg^{2+}$ ion is shown as light green sphere. The cryo-EM map is shown as blue or black mesh for MgADP and the protein, respectively, with a higher contour level for MgADP. Polar contacts are indicated by yellow dashes. **d,** The distance from the helical part of the R domain to M10 and M10' (shown as cartoon) is very similar, the assignment is based on a short extension at the N terminus of the R-helix that points towards M10. The cryo-EM map is shown as grey mesh, residues of the non-helical part of R and M10 are shown in stick representation.

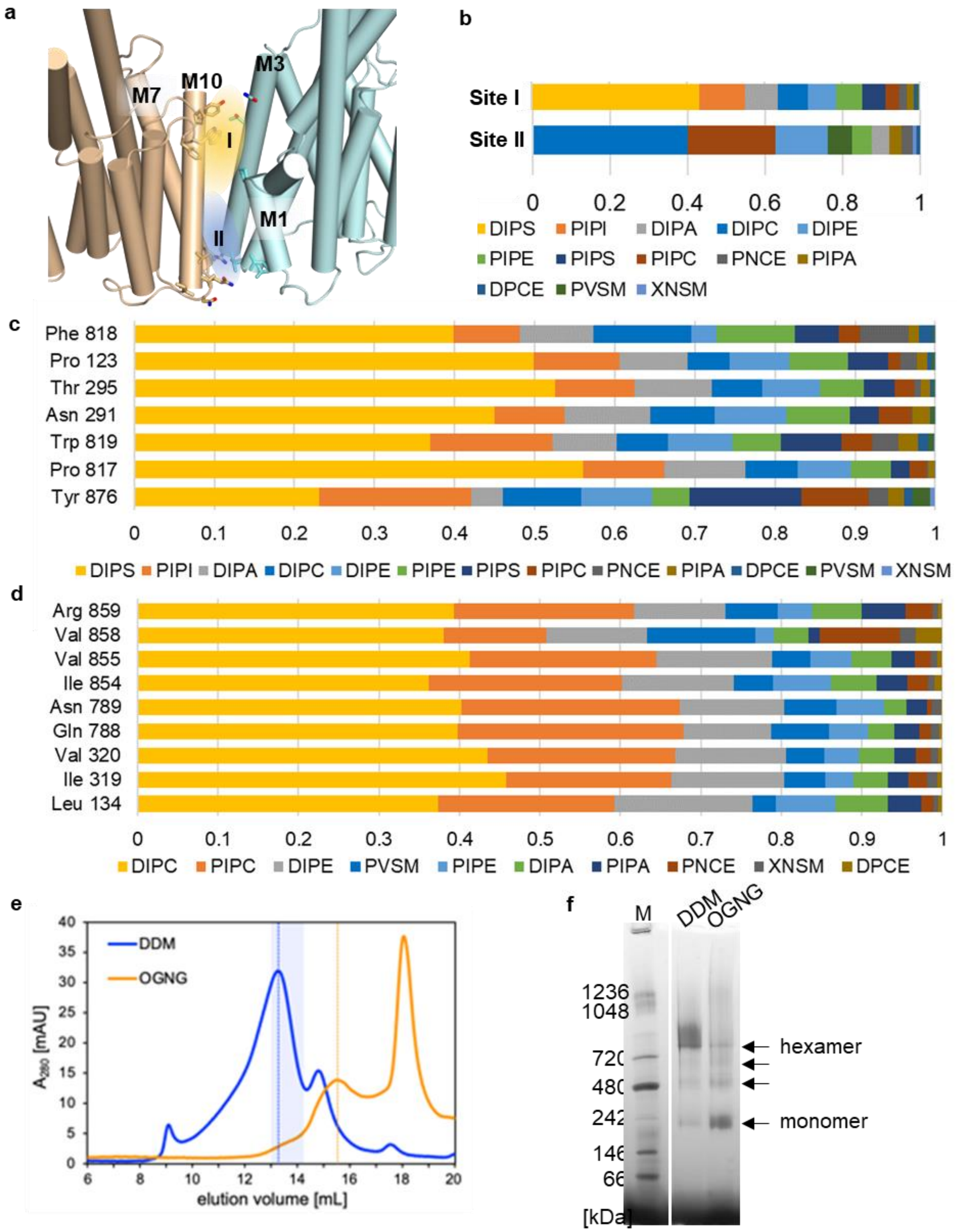

**Supplementary Figure S4 | Lipid binding and detergent effects on Pma1**

**a,** Putative lipid binding sites at the monomer interface, suggested by MD simulations. **b,** Fractional lipid interaction times in Site I and II, defined as the number of simulation frames in which a lipid is within 0.6 nm of a given site. **c,d**

Fractional interaction times of all lipids with Pma1 residues surrounding Site I (c) and Site II (d) **e,** Size-exclusion chromatography of Pma1 in DDM (blue; highlighted fractions used for cryo-EM) or OGNG (orange). **f,** Native PAGE (3-12 %) of Pma1 in DDM or OGNG. Unlabelled arrows indicate potential Pma1 dimers and tetramers. M: molecular weight marker.

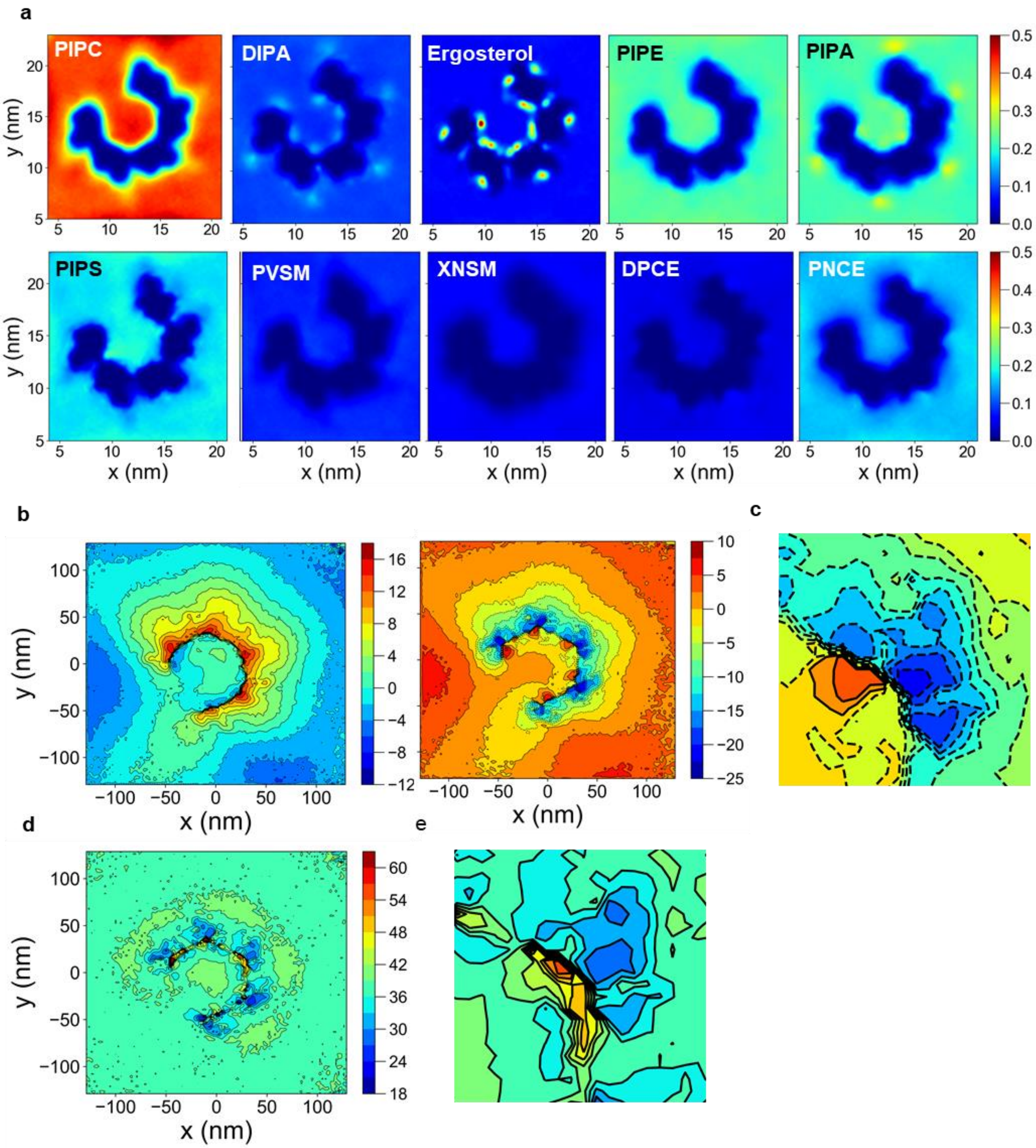

**Supplementary Figure S5 | Lipid density and membrane deformation maps**

**a,** Lipid density maps for all lipids used in the simulations besides DIPS and DIPC (see Fig. 4). Values correspond to average numbers of molecules per $nm^3$ and do not account for the respective membrane composition fraction. **b,** Average z-height position of lipid headgroup-phosphates in the outer leaflet (*left*) and the inner leaflet (*middle*). **c,** Zoom of **b** on the region of one Pma1 monomer. Values represent the z-height difference in Ångström relative to the value at coordinate -50, 110 (assumed to represent a membrane region unperturbed by protein or boundary effects). **d,** Average leaflet thickness between phosphates (inner leaflet minus outer leaflet) at each x, y coordinate. Scale is in Ångström. **e,** Zoom of **d** in the region of one monomer.

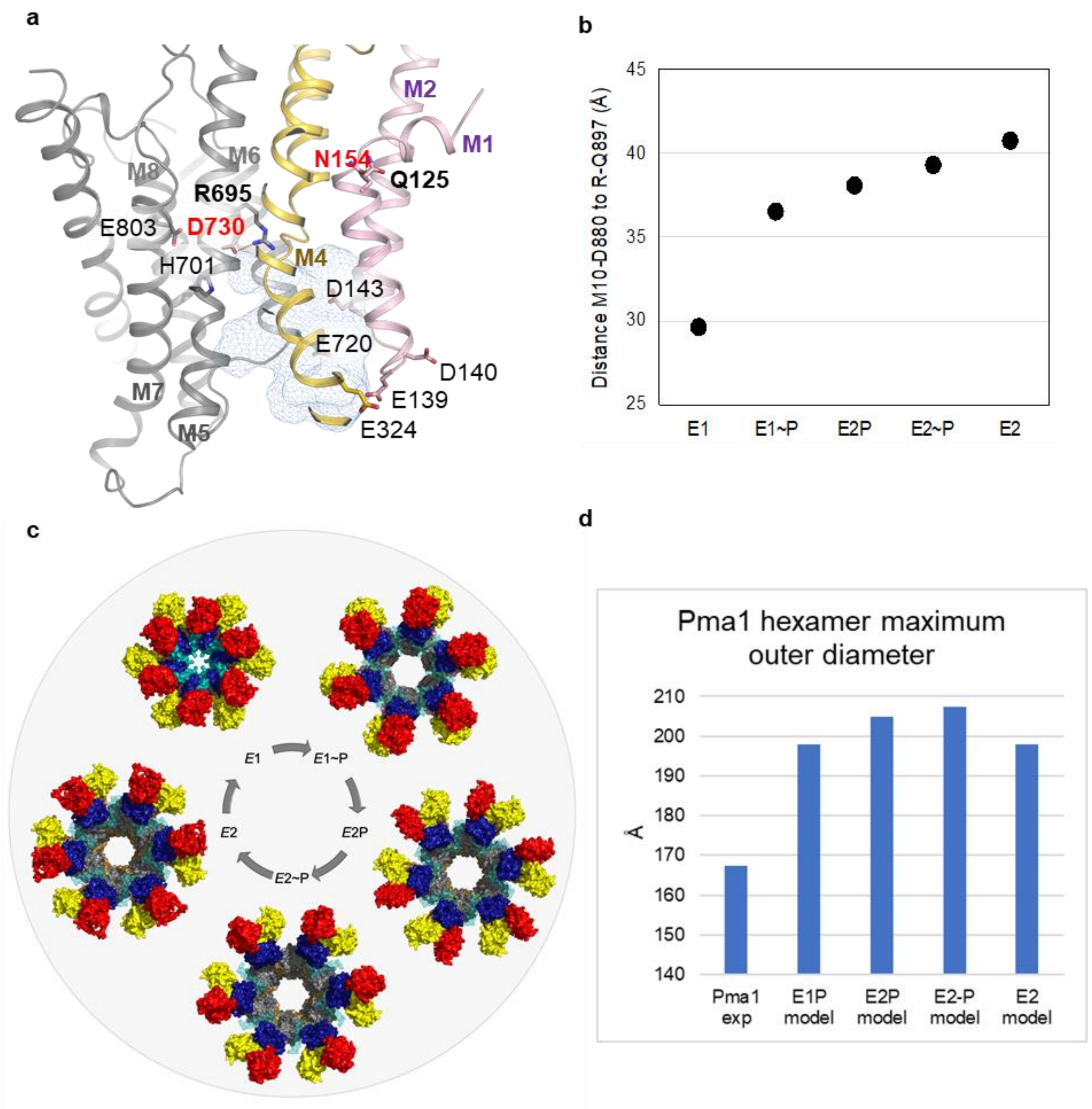

**Supplementary Fig S6 | Structural analyses of Pma1 homology models.**

**a,** Proton exit funnel between M1, M4 and M6 in the *E*2P homology model. Important residues for proton transport are shown as sticks. M1-2 are coloured pink, M3-4 gold and M5-10 grey. The proton acceptor/donor Asp730 (labelled in red) at the inner end of the funnel lies in bonding distance to Arg695, presuming a small side chain rotation of the latter (indicated bond: 4.9 Å). Asn154 (labelled in red), forms a putative hydrogen bond with Gln125 (2.8 Å). There is a cluster of negatively charged residues at the extracellular end of the exit funnel. **b,** Distance between the C-alpha atoms of the last residue of M10 (Asp880) and the first residue of the R-helix (Gln897) in the autoinhibited *E*1 structure and homology models, with the R-helix placed in its relative position to the P domain as in *E*1. **c,** Model of the catalytic cycle of hexameric Pma1. Homology models throughout the catalytic cycle were arranged into hexamers via alignment with M6-10 of the *E*1 hexamer. The R domain was placed in its relative position to the P domain as observed in *E*1. Nucleotide-binding (N) domain red, actuator (A) domain yellow, phosphorylation (P) domain blue, regulatory (R) domain cyan (transparent in homology models), M1-2 pink, M3-4 gold and M5-10 grey. **d,** Maximal outer diameter of the autoinhibited Pma1 *E*1 structure and homology models in states *E*1P, *E*2P, *E*2~P, and *E*2.

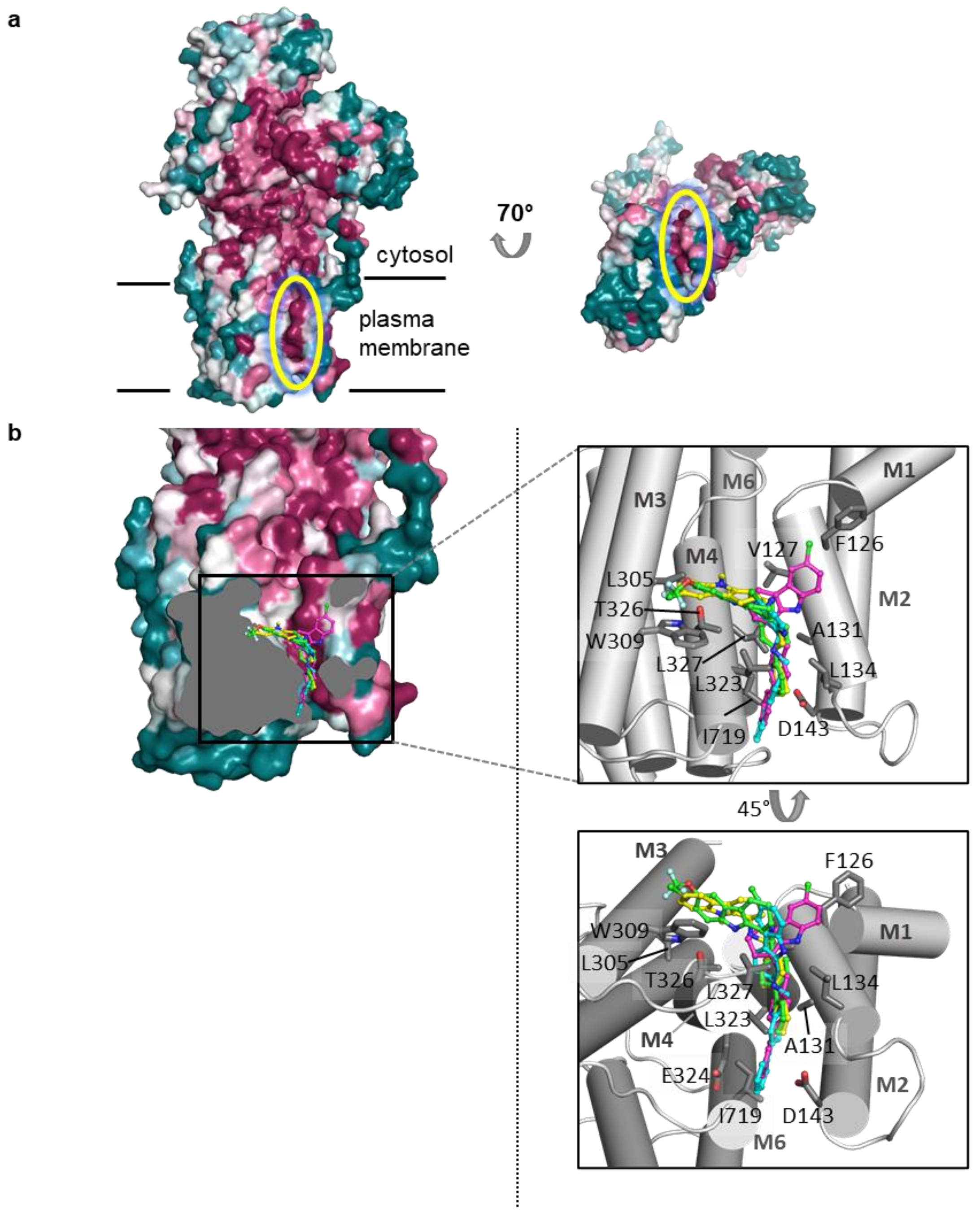

**Supplementary Fig. S7 | Compound docking into Pma1**

**a,** Conservation of Pma1 against proton pumps from human- and plant-pathogenic fungi (see species list in *Methods*). Calculated with ConSurf[45] and color-coded from purple (conserved) to blue-green (variable). Yellow circle highlights a conserved ridge along a putative drug-binding groove in the M domain, which extends into the extracytosolic side of Pma1.**b,** *Left*: Tetrahydrocarbazole compounds docked into Pma1 in the autoinhibited *E*1 state giving estimated affinities stronger than -9 kcal/mol (only one representative mode with the highest affinity score shown per compound): 9/S (green), 10/R (blue), 10/S (pink), 11/R (yellow). The protein surface is coloured as in **a**.

*Right:* Enlarged view of the putative inhibitor binding site with the protein shown as grey cartoon. Residues involved in binding of most compounds are shown as sticks.

### Supplementary Tables:

#### Supplementary Table S1 | Cryo-EM data collection, refinement and validation statistics

| | Hexamer<br>(EMDB-12644)<br>(PDB 7NY1) | | Monomer<br>(EMDB-12638)<br>(composite of EMDB-12642 /-12641 /-12643)<br>(PDB 7NXF) | |
|---|---|---|---|---|
| **Data collection and processing** | | | | |
| Magnification | 105’000 | | 105’000 | |
| Voltage (kV) | 300 | | 300 | |
| Electron exposure (e–/Å$^2$) | 42 | | 42 | |
| Defocus range (µm) | -1.3 to -2.5 | | -1.3 to -2.5 | |
| Pixel size (Å) | 0.837 | | 0.837 | |
| Symmetry imposed | C6 | | C1 | |
| Initial particle images (no.) | 483’742 | | 483’742 | |
| Final particle images (no.) | 59’511 | | composite (214’647* / 293’999* / 261’196*) | |
| Map resolution (Å) | 3.26 | | 3.11 | (3.25 / 3.17 / 3.15) |
| FSC threshold | 0.143 | | 0.143 | |
| **Refinement** | | | | |
| Initial model used (PDB code) | -/- | | -/- | |
| Model resolution (Å) | 3.35 | | 3.19 | |
| FSC threshold | 0.5 | | 0.5 | |
| Model resolution range (Å) | 318.06 – 3.35 | | 318.06 – 3.19 | |
| Map sharpening *B* factor (Å$^2$) | non-linearly postprocessed | | -26.79 | (not sharpened) |
| Model composition | | | | |
| Non-hydrogen atoms | 38430 | | 6405 | |
| Protein residues | 4974 | | 829 | |
| Ligands | K(6), Mg(6), ADP(6) | | K(1), Mg(1), ADP(1) | |
| *B* factors (Å$^2$) | | | | |
| Protein | 82.25 | (19.28 – 155.87) | 82.25 | (19.28 – 155.87) |
| Ligand | 107.30 | (61.94 – 109.49) | 107.30 | (61.94 – 109.49) |
| R.m.s. deviations | | | | |
| Bond lengths (Å) | 0.006 | | 0.006 | |
| Bond angles (°) | 0.810 | | 0.810 | |
| Validation | | | | |
| MolProbity score | 2.09 | | 1.96 | |
| Clashscore | 8.99 | | 12.46 | |
| Poor rotamers (%) | 0 | | 0 | |
| Ramachandran plot | | | | |
| Favored (%) | 92.1 | | 92.1 | |
| Allowed (%) | 7.9 | | 7.9 | |
| Disallowed (%) | 0 | | 0 | |

### Supplementary Table S2 | Residues involved in intra- and intermolecular hexamer contacts

| **Contacts involving the R domain** | | | | | |
|---|---|---|---|---|---|
| **R – P** | | **R – P’** | | **R – R’/ R” – R** | |
| Pro893 | Gly589 | Phe905 | Val562 | Leu902 | Ser892 |
| Lys 894 | Met592 | Leu909 | Gly563 | | Pro893 |
| Arg900 | Gly594 | Val912 | Arg566 | | |
| Glu903 | Ser595 | Ser913 | Asn577 | | |
| Asp904 | Tyr598 | Thr914 | Ile578 | | |
| Val907 | Asp599 | His916 | Tyr579 | | |
| Arg911 | Glu602 | Glu917 | Arg583 | | |
| | Arg625 | | Asp500 | | |
| | | | Phe600 | | |

| **Residues involved in the intermolecular contact within the M domain** | | |
|---|---|---|
| M3 / M4 | M7 / L7-8 | M10 |
| Thr295 | Ile772 | Ile862 |
| Ile299 | Thr775 | Phe863 |
| Ile302 | Thr776 | Cys869 |
| Leu306 | Gly784 | Ile870 |
| Trp309 | Gly785 | Tyr876 |
| Val310 | Ile786 | Ile877 |
| Phe313 | Gln788 | |
| Tyr314 | | |
| Pro318 | | |
| Ile319 | | |

**Supplementary Table S3 | Lipid composition used in the coarse-grained molecular dynamics simulations.**

| Lipid name | Head group | Tail | Net charge | Content in inner leaflet (%) | Content in outer leaflet (%) |
|---|---|---|---|---|---|
| PIPC | Phosphatidylcholine | C16:0/18:2 | 0 | 11 | 28 |
| DIPC | Phosphatidylcholine | di-C16:2-C18:2 | 0 | 5 | 14 |
| PIPE | Phosphatidylethanolamine | C16:0/18:2 | 0 | 8 | 13 |
| DIPE | Phosphatidylethanolamine | di-C16:2-C18:2 | 0 | 4 | 7 |
| PIPA | Phosphatidic acid | C16:0/18:2 | -2 | 11 | 11 |
| DIPA | Phosphatidic acid | di-C16:2-C18:2 | -2 | 5 | 5 |
| PIPS | Phosphatidylserine | C16:0/18:2 | -1 | 17 | 0 |
| DIPS | Phosphatidylserine | di-C16:2-C18:2 | -1 | 9 | 0 |
| PIPI | Phosphatidylinositol | C16:0/18:2 | -1 | 8 | 0 |
| XNSM | Sphingomyelin | C(d24:1/24:1) | 0 | 2 | 2 |
| PVSM | Sphingomyelin | C(d18:1/18:1) | 0 | 4 | 4 |
| DPCE | Ceramide | C(d18:1/18:0) | 0 | 3 | 3 |
| PNCE | Ceramide | C(d18:1/24:1) | 0 | 8 | 8 |
| ERGO | Ergosterol | – | 0 | 5 | 5 |
